# Morphometry and diversity of birds in cocoa agroforestry and silvopastoral systems in the Colombian Amazon

**DOI:** 10.1101/2025.01.28.635237

**Authors:** Elkin Damián Reyes-Ramírez, Jenniffer Tatiana Díaz-Cháux, Alexander Velasquez-Valencia

**Author notes:** Corresponding author, (JTD).

## Abstract

The primary cause of deforestation in the Amazon region is extensive traditional cattle ranching, which is considered an important economic activity in several departments of the region, particularly in Caquetá. This research aimed to determine the influence of vegetation cover on the morphometry and diversity of birds in agroforestry systems with cocoa and silvopastoral practices in the Colombian Amazon. Sampling was conducted in eight locations in the Caquetá department using mist nets between January and November 2023. In each location, five effective sampling days were carried out, and for each captured individual, weight and morphometric measurements of the bill, wings, legs, and tail were recorded. Based on the wing measurements, the Kipp’s index was calculated, relating the morphology to the dispersal capacity of the birds. A total of 350 individual birds were recorded, distributed across 77 species and 20 families in the sampled agroforestry systems with cocoa and silvopastoral practices. The Early Brush cover type exhibited the highest richness and abundance, and it was determined that the variation in the morphometric traits of the birds is associated with the type of habitat. The species accumulation curve from the collected mosaics allowed for the documentation and analysis of community richness in these two systems. It was determined that agroforestry systems with cocoa and silvopastoral practices, due to their tree structures simulating a heterogeneous habitat, have the capacity to host many species and are therefore important for the conservation of avifauna. Anthropization and fragmentation of natural habitats exert pressures on birds, leading to modifications in their morphometric traits to adapt to their environmental conditions.

## Introduction

In the last decade, the Colombian Amazon territory has lost 1 858 285 natural forest hectares, and one of the deforestation hot spots is in Caquetá (1). In this sense, the principal reason for the habitat loss is the extensive livestock (2). Clearly, livestock is considered the principal economic activity in the department (3). For this reason, this activity has presented a rapid increase over the years. As a result, this increase has caused the deforestation of more than 3 585 forest hectares in 2024 (4).

Thus, deforestation directly affects the composition and structure of the communities as floristics as fauna (5). Likewise, it generates the loss of the ecosystemic services of these communities (6). In that regard, birds are one of the taxonomic groups susceptible to the degradation of the habitat and the change in land use (7–10). Hence, the transformation of the natural landscape allows the loss of connectivity between habitat fragments, which affects the birds’ species’ mobility (11). This condition influences the reproductive and forager behavior of birdlife since the structural complexity of vegetation determines the ecological and functional diversity of birds (12).

As a mitigation mechanism for the deforestation effects, in Caquetá, eco-friendly production systems are being implemented to preserve biodiversity (6, 7). Therefore, by implementing cocoa agroforestry systems (SAFc) and silvopastoral systems (SSP), the goal is to manage rural landscapes through efficient agricultural and livestock production systems (15). These systems are distinguished by their high density of shrubs and trees, creating a diverse habitat that provides resources for various bird species (16).

Incorporating trees as an integral part of the SSP significantly enhances livestock production systems, reduces the environmental impact of conventional livestock practices, boosts animal welfare, and leads to higher productivity levels (10, 11). On this matter, SSP and SAFc improve the ecological structure and connectivity of the landscape and conserve biodiversity and its ecological functions (19). Therefore, SSP and SAFc are crucial in conserving bird species in fragmented landscapes by offering shelter, resting places, nests, and sustenance (20).

In this context, alterations in the makeup of bird populations within fragmented habitats are observable (7). Moreover, landscape transformation’s effect on these communities’ structures has been documented. Hence, the foraging guilds have studied these changes (18, 19). Certain guilds are advantaged when landscapes are developed with varied covers and more superficial plant structures (23). The pattern is evident in birds that feed on herbaceous seeds or flying insects, primarily found in open spaces (24). Nonetheless, these changes affect forest-dependent species, such as those that consume fruits, insects, or undergrowth (25–28).

In the same line of thought, some investigations pretend to explain these effects through functional groups. These investigations include the analysis of birds’ morphometrical variations in different ecological conditions (8, 22, 23). According to Jackson & Fahrig (31) and Matyjasiak et al. (32), the separation of these areas has led to alterations in the wing shapes and sizes of migratory species.

Similarly, it has been shown that the quality of patches and vegetation cover types in fragmented landscapes influences the body state and birds’ morphometric conditions (33). Based on the information above, this study assessed how cocoa agroforestry production systems and silvopastoral impact bird diversity and morphometrics in the Colombian Amazon. Likewise, the ability of bird species to disperse and move across the landscapes within productive systems was evaluated.

## Materials and methods

### Study area

This investigation was carried out in eight localities with sustainable production systems in Caquetá, in the northwestern region of the Colombian Amazon (Fig. 1). The area has an average annual temperature of 25.6 °C and an average annual precipitation between 2 400 and 3 422 mm. The relative humidity is between 69 % and 86 %; June and July have the highest values, and the year’s first quarter has the lowest (34).

**Fig 1.**
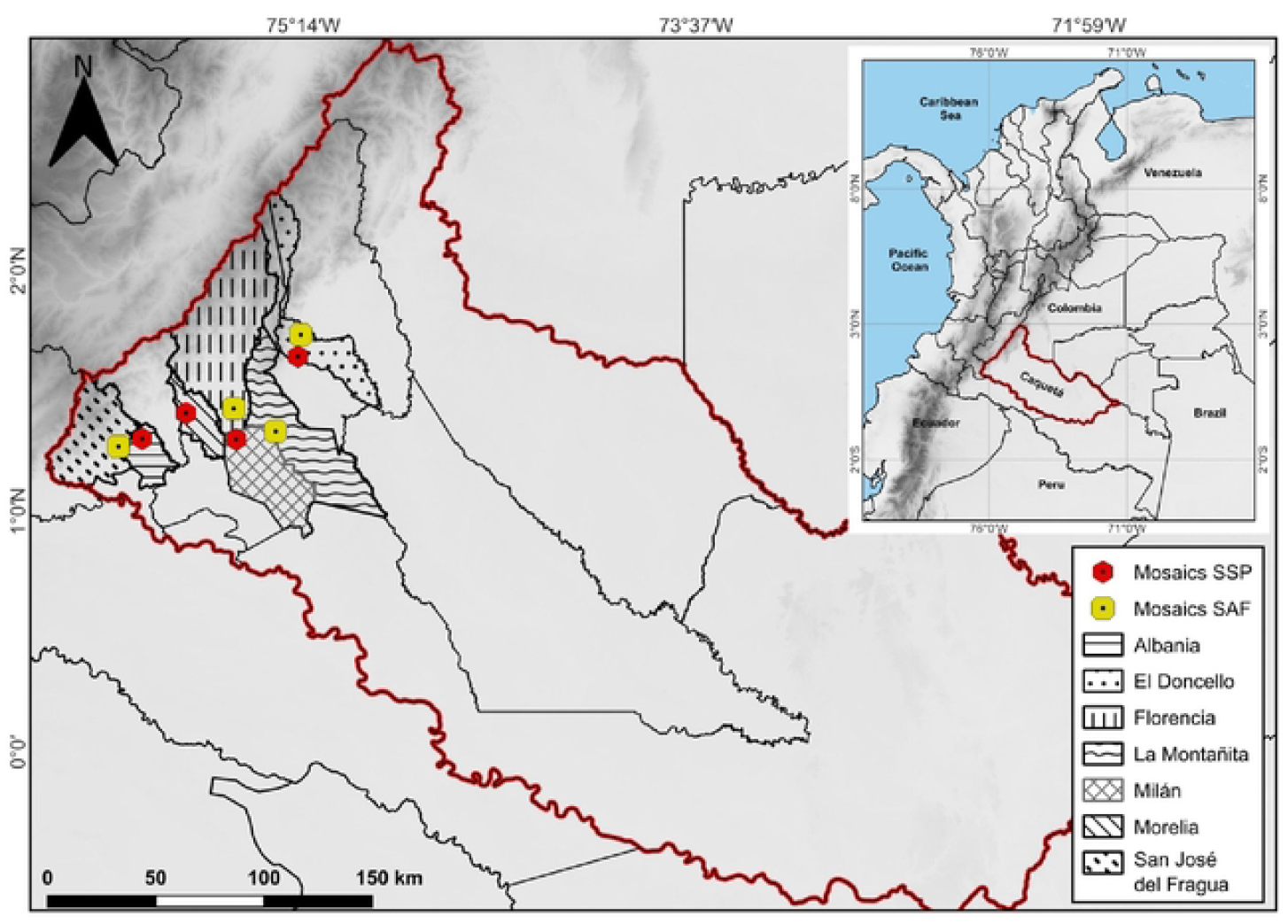
Location of landscape mosaics with agroforestry and silvopastoral systems in the Colombian Amazon. SAFc: TR (El Triunfo – Doncello), BA (Batalla 13 – Florencia), SR (Santa Rosa – San José del Fragua), TE (El Tesoro – Montañita). SSP: PO (El Porvenir – Albania), VE (La Vega – Doncello), VM (Villa Mery – Morelia), ES (Esmeraldas - Milán).

In each locality, one km^2^ mosaic was established as a midpoint within the landscapes of agroforestry and silvopasture systems. In the mosaics, a grid composed of five linear transects separated by 0.25 km was established. The classification of vegetation covers was developed based on the CORINE Land Cover methodology adapted by Velasquez Valencia and Bonilla-Gómez (3, 14).

Therefore, seven covers were classified in every sampled mosaic. First; cocoa agroforestry crops (CAC) that are constituted for planting *Theobroma cacao* (L.) combined with trees or shrub species. Second; Paddocks with scattered trees (PAD) are characterized by the presence of trees taller than five meters distributed in a dispersed manner. Thirdly; paddocks that have been weeded (PEN) feature a covering of *Brachiaria* sp. grasses, with weed heights not exceeding 1.5 meters. Likewise, the hollow paddocks s (PPH) are pastures located in flood zones between the valleys of the hills.

Furthermore, clean paddocks (PPL) consist of areas covered by pastures, predominantly featuring *Brachiaria* sp. in over 70 % of the land, in contrast to the early stubble (RTT) areas, which exhibit vegetation of a shrubby-herbaceous nature, standing between two and five meters tall, with a dominant diameter at breast height of less than 10 cm. In conclusion, areas classified as old stubble (RTV) consist of vegetation, predominantly trees, that are over ten years old, reaching heights of 8 meters, with a dominant diameter at breast height greater than 10 cm, and featuring an uneven canopy.

### Birds sampling

The diversity of birds was assessed using mist nets across transects in mosaic patterns from January to November 2023. Nine mist nets were installed within a total of 90 meters per transect; this design made it possible to intercept a greater mobility path of the birds between the vegetal covers (36–38).

For five efficient days, mosaics conducted the bird census, with the mist nets being unfolded from 6:00 hrs to 11:00 hrs, a span of five hours that captures the peak activity period of the birds, ensuring checks were made every 20 minutes. Each captured individual was measured to register the morphometric traits of beaks, wings, legs, and tails. In examining the traits of the beak, measurements included beak height (ALT), total culmination (CTO), exposed culmen (CEX), and commissure (COM). In examining the traits of wings, the following characteristics were observed: extended wing (AEX), flat wing (APL), primary feathers (PRI), and secondary feathers (SEC).

Moreover, measures were taken for the tarsus length (LTA), tail length (LCO), total body length (LTO), and weight of the individual. Furthermore, the taxonomical determination of species was carried out following the nomenclature of the South American Classification Committee (SACC) (39). Besides, the verification of the species was performed by relying on the Colombian birds’ guide (18, 19) and the official list of Colombian birds in 2022 (42).

The Kipp’s Index (43) was calculated based on wing morphometric traits, including the length from the first secondary feather to the tip of the largest primary feather, measured with the wing totally folded (44). This index relates the morphology of the wing to birds’ capacity to disperse (45) and migrate.

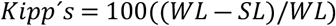

Where WL is the total length of the closed wing (mm) and SL is the distance from the carpus to the tip of the first secondary feather (mm) (46).

### Data analysis

The birds’ diversity was analyzed using the index’s calculation of diversity through the program Past—Palaeontological Statistics, version 1.81 (47). The curve of species accumulation was built through the program EstimateS version 9.1 Windows (48), and the Jaccard similarity index was estimated to evaluate the similarity of species between vegetation covers.

The morphometric traits were assessed using Analysis of Variance (ANOVA) with the software Infostat version 2.0 (49). Tukey’s method was used to assess the impact of the productive systems (SAFc and SSP) on the bird species and the morphometric traits. To establish the relationship between the morphometric traits of bird species and the production systems, an ANOVA was used with the Infostat version 2.0 program (49) using Tukey’s method, considering the morphometric traits per species captured in each system.

## Results

### Composition and structure of birds’ community in vegetation covers

A total of 82 transects were sampled; in every transect, 90 meters of nets were installed, which is a total of 185 hours/net per mosaic. 350 individual birds were captured, representing 77 species, 20 families, and eight orders in the agroforestry systems with cocoa (SAFc) and silvopastoral (SSP) mosaics sampled. SAFc and SSP shared four covers (PEN, PPL, RTT, RTV). LAG and RIO were only present in the SSP, and CAC only in SAFc.

The most collected species were *Ramphocelus carbo* (45 individuals) and *Chionomesa fimbriata* (31 individuals); 29 % of the species presented a one-and-only record. In SAFc, 51 species and 168 individuals were collected, while in SSP, 52 species and 182 individuals were collected. Neither were differences found in the wealth among the assessed systems (KW=1.16; n=82; P> 0.05; KW= 2; n=82; P> 0.05) nor in the covers (KW=10.24 n=82; P> 0.05; KW= 10.59; n=82; P> 0.05).

The RTT and PPL covers showed the highest wealth, while RTV and PPH had the lowest values. PPH had the highest dominance, followed by RTV. The highest evenness (1-D) was found in PPL and RTT, and the highest diversity was observed in the RTT and PPL coverages (Table 1).

**Table 1.**
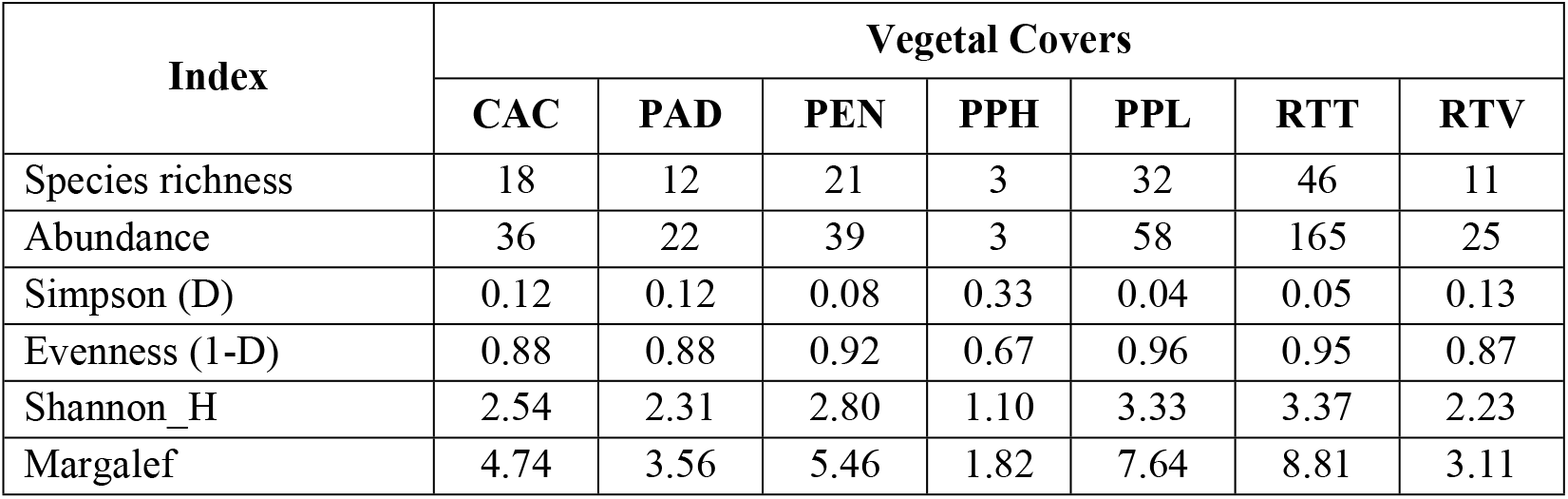
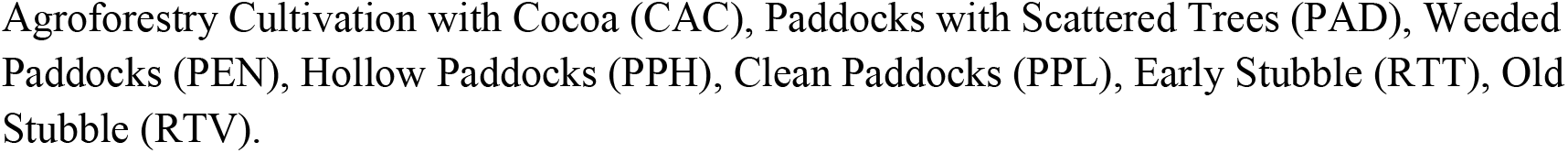
Diversity indices of the community of birds captured in the vegetal covers classified in cocoa agroforestry and silvopastoral systems of the Amazon, Colombia.

| Index | Vegetal Covers |  |  |  |  |  |  |
| --- | --- | --- | --- | --- | --- | --- | --- |
|  | CAC | PAD | PEN | PPH | PPL | RTT | RTV |
| Species richness | 18 | 12 | 21 | 3 | 32 | 46 | 11 |
| Abundance | 36 | 22 | 39 | 3 | 58 | 165 | 25 |
| Simpson (D) | 0.12 | 0.12 | 0.08 | 0.33 | 0.04 | 0.05 | 0.13 |
| Evenness (1-D) | 0.88 | 0.88 | 0.92 | 0.67 | 0.96 | 0.95 | 0.87 |
| Shannon_H | 2.54 | 2.31 | 2.80 | 1.10 | 3.33 | 3.37 | 2.23 |
| Margalef | 4.74 | 3.56 | 5.46 | 1.82 | 7.64 | 8.81 | 3.11 |

Agroforestry Cultivation with Cocoa (CAC), Paddocks with Scattered Trees (PAD), Weeded Paddocks (PEN), Hollow Paddocks (PPH), Clean Paddocks (PPL), Early Stubble (RTT), Old Stubble (RTV).

Additionally, 87 % of the expected species were collected during sampling, as indicated by the non-parametric indices of Chao 1 and Jacknife 1 (Fig. 2).

**Fig. 2.**
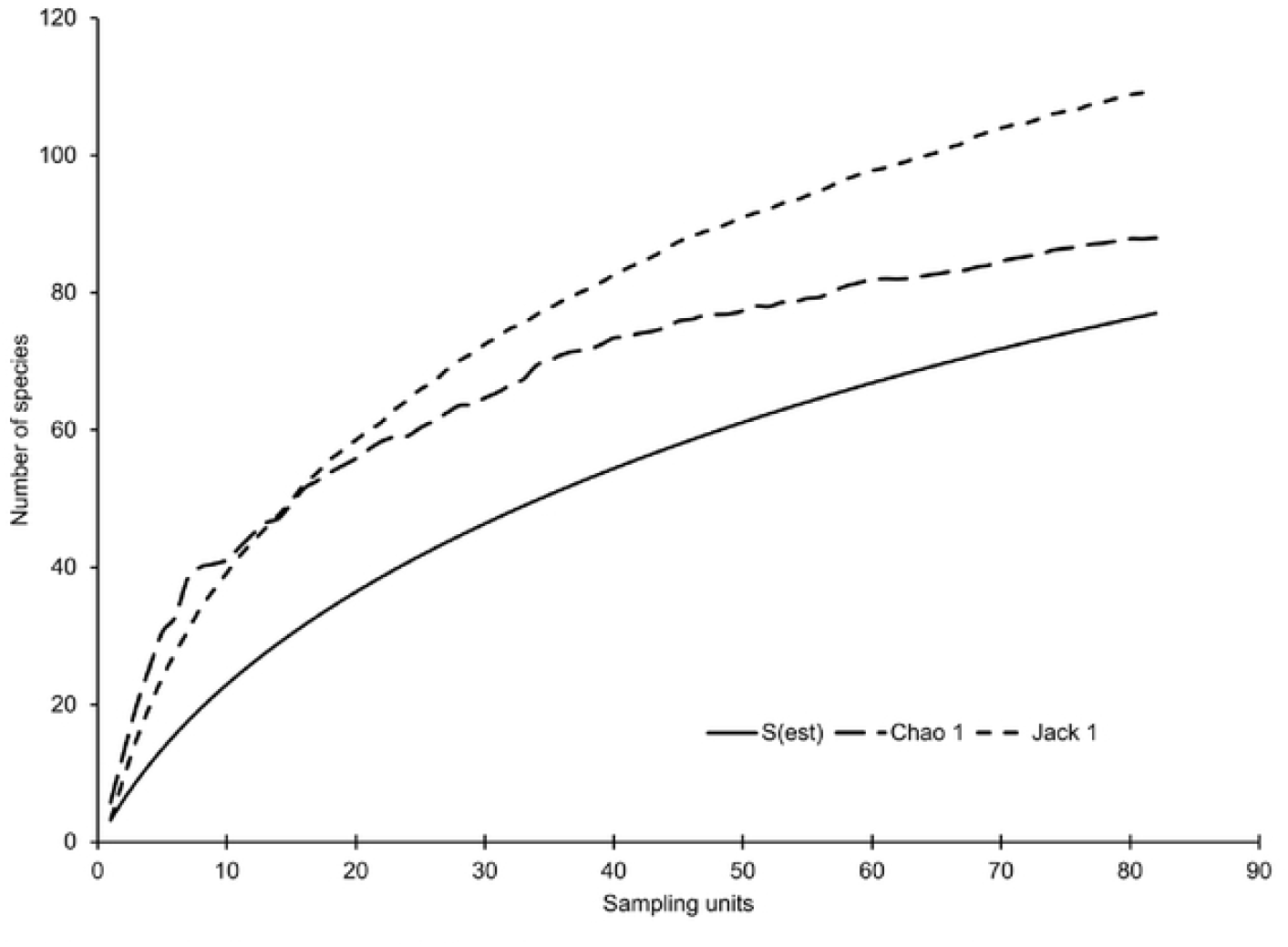
Accumulation curve of bird species captured in agroforestry and silvopastoral systems in the Amazon, Colombia.

Additionally, twenty-six species were registered in both systems SAFc and SSP, species such as *Camptostoma obsoletum, Columbina talpacoti, Glyphorynchus spirurus, Manacus manacus, Schistochlamys melanopis*. The covers with the highest similarity in species composition were RTT and PPL, with 15 shared species, followed by the covers CAC and PEN. Thus, it is vital to mention that the covers PPH, PAD, and RIO presented the lowest similarity compared to the other covers (Fig. 3). The PPL Clean Paddocks presented the most significant number of exclusive species such as *Dryobates passerinus, Machaeropterus striolatus, Sporophila murallae*, and *Nyctidromus albicollis* (Table 2). Followed by RTT coverage with exclusive species such as *Capito aurovirens, Glaucis aeneus, Pteroglossus inscriptus, Rupornis magnirostris*. Four hedges shared more than ten species, and the rest less than five. In total, 31 species are found in more than one location. These include *Chionomesa fimbriata, Ramphocelus carbo, Sporophila angolensis, Phaethornis hispidus, and Hemitriccus Zosterops*. Among these species, three were recorded in six covers, five in at least four covers, and eight species in at least three covers.

**Table 2.** Distribution of richness species in vegetation covers classified in cocoa agroforestry and silvopastoral systems of the Amazon, Colombia. Exclusive species on the diagonal, shared species below the diagonal, and percentage similarity above the diagonal

|  | CAC | LAG | PAD | PEN | PPL | RIO | RTT | RTV |
| --- | --- | --- | --- | --- | --- | --- | --- | --- |
| CAC | **4.00** | 5.00 | 5.00 | 16.13 | 20.41 | 5.26 | 20.41 | 17.86 |
| LAG | 1.00 | **2.00** | - | 10.53 | 4.76 | 25.00 | 4.76 | 5.88 |
| PAD | 1.00 | - | **1.00** | 5.00 | 2.33 | - | - | 5.88 |
| PEN | 5.00 | 2.00 | 1.00 | **3.00** | 22.92 | 5.26 | 22.92 | 22.22 |
| PPL | 10.00 | 2.00 | 1.00 | 11.00 | **15.00** | 2.38 | 34.43 | 16.67 |
| RIO | 1.00 | 1.00 | - | 1.00 | 1.00 | **1.00** | 2.38 | 6.25 |
| RTT | 10.00 | 2.00 | - | 11.00 | 21.00 | 1.00 | **14.00** | 19.15 |
| RTV | 5.00 | 2.00 | 1.00 | 6.00 | 8.00 | 1.00 | 9.00 | **3.00** |

Agroforestry Cultivation with Cocoa (CAC), Paddocks with Scattered Trees (PAD), Weeded Paddocks (PEN), Hollow Paddocks (PPH), Clean Paddocks (PPL), Early Stubble (RTT), Old Stubble (RTV).

**Fig. 3.**
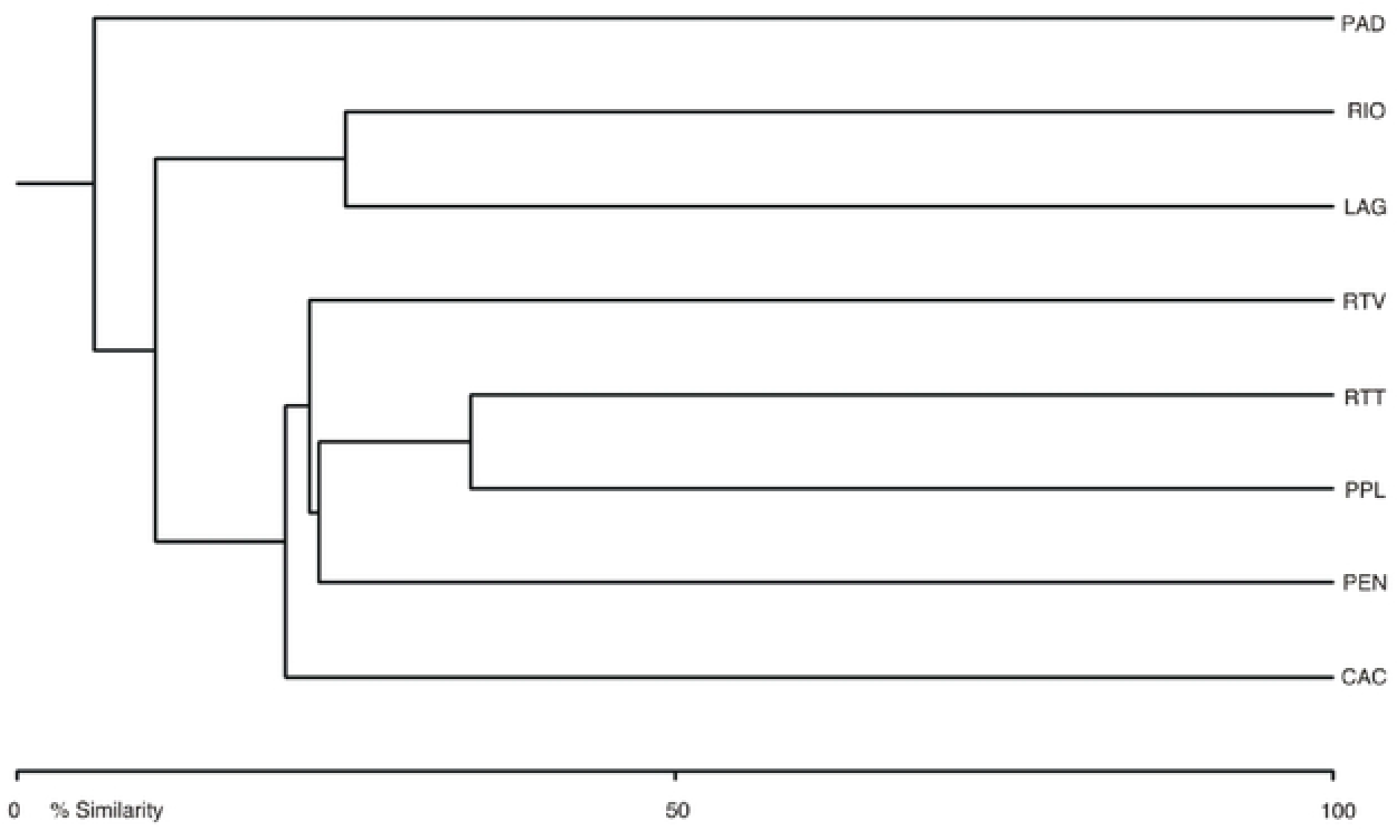
Jaccard’s similarity index of bird species by vegetation cover in landscape mosaics with agroforestry and silvopastoral systems of the Amazon, Colombia.

### Relation of the morphometric traits of birds’ diversity in the cocoa agroforestry and silvopastoral systems

Only three species presented differences in the Kipp’s index’s values among the systems *Arremonops conirostris* (F=7.99; gl=10; P<0.05); *Chionomesa fimbriata* (F=6.39; gl=30; P<0.05), and *Ramphocelus carbo* (F=4.86; gl=44; P<0.05). In the three species, this index presented its highest value in SAFc.

Only eight morphometric variables presented differences among the systems SAFc and SSP in seven species. In the species *Phaethornis hispidus*, the average values of LTA (F=6.82; gl=7; P<0.05) and PES (F=6.58; gl=7; P<0.05) were higher in SSP. Equally, the LTA measures in the species *Ramphocelus carbo* (F=73.44; gl=43; P<0.05) and PRI in the species *Hemitriccus zosterops* (F=5.77; gl=9; P<0.05) and *Leptopogon amaurocephalus* (F=7.17; gl=7; P<0.05) presented higher values in SSP. While in SAFc the morphometric variables CEX, CTO, LTA, AEX, APL, and ALT were higher in the species *Arremonops conirostris* CTO (F=7.37; gl=10; P<0.05) and CEX (F=37.29; gl=10; P<0.05); *Thraupis episcopus* LTA (F=63.48; gl=15; P<0.05); *Sporophila angolensis* has distinctive traits in both of its wings, known as AEX (F=5.11; gl=15; P<0.05) and APL (F=5.18; gl=15; P<0.05); and *Thraupis episcopus* ALT (F=19.07; gl=4; P<0.05).

## Discussion

The accumulation curve of the collected species in the agroforestry and silvopastoral systems in the sampled mosaics allows for documenting and analyzing the wealth of the communities within both systems. Although the wealth of the average collected species in the mosaics of SSP and SAF systems did not differ, it was higher in the silvopastoral systems. These results are similar to those found by Cruz-Trujillo (50), where no significant differences were observed in these two attributes across the production systems. Nevertheless, they register a higher number of species in the agroforestry systems. In the same line, Velásquez-Valencia and Bonilla-Gomez (7) discovered that agroforestry crops and systems with natural regeneration associated with secondary vegetation had higher wealth. It is important to note that in this study, species were recorded using observation points.

The most common species was *R. carbo*, found in seven out of eight sampled locations. This species comprehends generalist habits (51). That is why it does not have any preference towards a specific habitat, and it can be found in anthropized environments, so *R. carbo* has adaptability and plasticity capacities (52). These results are similar to some investigations in Caquetá, where the Thraupidae family presented a high abundance of SAF and SSP (50, 53).The early stubble cover presented the highest bird diversity, possibly related to its high dominance in the sampled mosaics. It is essential to emphasize that the presence of vegetation patches allows the habitation of typical birds from open areas or those tolerant to transformation. This is why these kinds of covers are appropriate habitats for birds (53, 54).

Given the differences that silvopastoral and agroforestry systems may have in terms of their composition and tree structure, the diversity of birds may vary in each system (7). However, they are of great importance for the conservation of avifauna (56–58). According to Giraldo et al. (53), the great diversity of birds that silvopastoral systems house is due to the shelter and food conditions it provides, such as insects and fruits, unlike areas such as clean pastures that are characterized by not having tree cover. Besides, most birds do not find sufficient resources for their stability, and ecologically, these individuals tend to prefer other habitats (59–61). Likewise, the organization of agroforestry systems generates different habitats that facilitate the regulation of climate and temperature, and they are also essential refuge sites for biodiversity, especially for birds (62).

The morphometric traits of birds provide valuable information about the adaptations these individuals present based on their environment and the functions they fulfill in an ecosystem (63). The analysis of Kipp’s index showed higher values for SAFc and lower values for SSP; this is related to the specialization of the bird. Generalist species found in different vegetation covers have great flight capacity. Therefore, their wings are enormous (64). These changes in bird wing morphology confirm that habitat type exerts pressure, leading to trait modifications over time (32, 58).

Kiat and O’Connor (66) establish that the form of birds’ wings is tightly related to mobility and habitat type. Subsequently, Revelo-Hernández et al. (67) corroborated that there is a direct relation between the characteristics of the habitat and the avian morphology, confirming that habitat transformations have the capacity to change morphological traits in bird communities. However, the results of this investigation are different (67) since these authors found that birds from silvopastoral systems present longer wings, which indicates that in these systems, the degree of disturbance is the determining factor that puts pressure on the morphometry of these individuals.

The differences found between the form and size of the *Arremonops conirostris* beak are directly related to this species’ foraging behavior and feeding habits (61, 62). This species exhibits its highest values of these traits in the SAFc systems, where more complex vegetation covers are associated with a broader diversity of fruits, shrubs, and trees (70). Therefore, this investigation allows perceiving a change in the beak size of this species, possibly to maximize the foraging in the existent resources of SAFc. Besides, these differences allow birds to inhabit the same areas without competing for food. Disturbances and anthropization of natural habitats mean that biodiversity, especially birds, are isolated in areas with different ecological conditions. In this scenario, only species that can tolerate new habitat conditions respond positively (64, 65). This includes species that start to change their physical characteristics in response to the challenges of new habitats, as demonstrated in this research.

Species reported as specialists, with habitats limited to dense forests, have undergone morphological changes, enabling them to colonize other ecological niches (73). This confirms that the variation in the tarsus length in *Phaethornis hispidus* is present because of the structural conditions of SAFc and SPP. In the silvopastoral systems dominated by a pasture matrix, the edge effect on the patches of remaining vegetation includes a greater incidence of winds inside the covers. This condition could affect this species’ perching ability, facilitated by larger tarsi. However, more data must be obtained from this species to test this hypothesis. Consequently, Eliason et al. (74) state that this morphometric variable is lower in birds from forested areas as opposed to those that inhabit open regions such as SSP. However, the results of this research differ from Revelo Hernández et al. (67), who found that tarsi length is more remarkable in SSP. Although they attribute these changes to the adaptations of birds to the type and quality of habitat. Likewise, variations in body weight can be influenced by ecological conditions such as habitat quality, food availability, temperature, flight capacity, and dispersion (75–77).

## Conclusions

The silvopastoral systems contribute to a higher cover diversity, including early stubble, where the highest birds’ diversity was captured. Moreover, Kipp’s Index showed how generalist species presented higher flying capacity in habitats such as SAFc, indicating that the wing size is much larger. This means that environmental pressures condition changes in allometry. Finally, the morphometric traits that presented significative differences per system indicate how birds adapt to their habitats through their morphometry in traits such as wings, beaks, or tarsus.

## Acknowledgments

This paper is part of the PhD investigation by Díaz-Cháux, J.T. in the sustainable development and natural sciences PhD at Universidad de la Amazonia. It comprehends a PhD excellence grand from Bicentenario conferred by Minciencias Colombia (code BPIN 2021000100036). The authors are grateful with Centro INBIANAM for facilitating the equipment for the field work. To Comité Departamental de Ganaderos del Caquetá and Asociación Departamental de Cultivadores de Cacao y Especies Maderables del Caquetá – ACAMAFRUT for facilitating the access to the fields. To Alejandro Navarro for constructing the thematic map of covers. To Mauren Andrea Ordoñez and Semillero de Investigación en Biodiversidad y Servicios Ecosistémicos -BySE for their support in the application and data collection phases.

## Author Contributions

**Conceptualization**: Elkin Damian Reyes Ramirez, Jenniffer Tatiana Díaz-Cháux.

**Data curation**: Elkin Damian Reyes Ramirez, Jenniffer Tatiana Díaz-Cháux, Alexander Velasquez Valencia.

**Funding acquisition**: Jenniffer Tatiana Díaz-Cháux, Alexander Velasquez Valencia.

**Investigation**: Elkin Damian Reyes Ramirez, Jenniffer Tatiana Díaz-Cháux.

**Resources**: Elkin Damian Reyes Ramirez, Jenniffer Tatiana Díaz-Cháux, Alexander Velasquez Valencia.

**Supervision**: Jenniffer Tatiana Díaz-Cháux, Alexander Velasquez Valencia.

**Validation**: Alexander Velasquez Valencia

**Writing – original draft**: Elkin Damian Reyes Ramirez

**Writing – review & editing**: Jenniffer Tatiana Díaz-Cháux, Alexander Velasquez Valencia

## Notes

### Competing Interest Statement

The authors have declared no competing interest.

